# Metabolic Processing of Selenium-based Bioisostere of *meso*-diaminopimelic Acid in Live Bacteria

**DOI:** 10.1101/2021.12.18.472960

**Authors:** Alexis J. Apostolos, Thameez M. Koyasseril-Yehiya, Carolina Santamaria, José Rogério A. Silva, Jerônimo Lameira, M. Cláudio N. Alves, M. Sloan Siegrist, Marcos M. Pires

## Abstract

The bacterial cell wall supports cell shape and prevents lysis due to internal turgor pressure. A primary component of all known bacterial cell walls is the peptidoglycan (PG) layer, which is comprised of repeating units of sugars connected to short and unusual peptides. The various steps within PG biosynthesis are often the target of antibiotics as they are essential for cellular growth and survival. Synthetic mimics of PG have proven to be indispensable tools to study bacterial cell growth and remodeling. Yet, a common component of PG, *meso*-diaminopimelic acid (*m*-DAP) at the third position of the stem peptide, remains challenging to build synthetically and is not commercially available. Here, we describe the synthesis and metabolic processing of a selenium-based bioisostere of a *m*-DAP analogue, selenolanthionine. We show that selenolanthionine is installed within the PG of live bacteria by the native cell wall crosslinking machinery in several mycobacteria species. We envision that this probe will supplement the current methods available for investigating PG crosslinking in *m*-DAP containing organisms.

## Introduction

The bacterial cell envelope is highly complex and its overall architecture can differ among the different classifications of bacteria (e.g., Gram-positive, Gram-negative, and mycobacteria) however, all possess a peptidoglycan (PG) layer. PG is a large biomacromolecule that encompasses the inner membrane of bacteria.^1^ It is essential for the viability of bacterial cells; a feature that is tied to the structural support it provides by maintaining cellular shape and rigidity. For almost all known bacteria, PG is made up of a backbone glycan strand connected to short, and highly unusual, peptides (often referred to as stem peptides). While there can be variability in the primary sequence of the stem peptide, some features remain constant. Stem peptides are initially biosynthesized as pentamers and most often contain the sequence L-Ala-D-iGlu-X-D-Ala-D-Ala, whereby X can be *meso*-diaminopimelic acid (*m*-DAP) or L-Lys (typically modified with a crossbridge).

The most varied position within the stem peptide is the third amino acid.^*1*^ Bacteria whose PG contain *m*-DAP (**Fig. 1A**) are diverse, ranging across Gram-positive bacteria such as *Bacillus subtilis* (*B. subtilis*) and *Lactobacillus plantarum* (*L. plantarum)*, Gram-negative bacteria such as *Escherichia coli* (*E. coli*) and *Pseudomonas aeruginosa* (*P. aeruginosa*), and mycobacteria such as *Mycobacterium tuberculosis* (*M. tuberculosis*).^*2, 3*^ Many of these bacteria are important for human health as these organisms can be dangerous pathogens and/or part of the microbiota community that colonize humans. The difference between *m*-DAP and L-Lys may be structurally subtle (a carboxyl on the ε carbon) but this variation can impose a significant impact on how PG fragments are sensed or metabolized.^*4-6*^ For instance, fragments of PG and other macromolecules released by bacterial cells are recognized by the human immune system as pathogen-associated molecular patterns (PAMPs), which are biomolecules that signal the presence of bacterial cells (a potential danger signal).^*7*^ A prominent PAMP receptor in human immune cells, NOD1, is activated by the minimal PG motif, D-iGlu-*m*-DAP (iE-DAP).^*8*^ However, the analogous PG fragment from L-Lys containing bacteria, D-iGlu-L-Lys, does not activate NOD1. Moreover, recent studies have demonstrated that amidation of *m*-DAP can greatly attenuate NOD1 activation, which suggests a potential pathway for *m*-DAP containing bacteria to remodel the PG layer to reduce recognition by the host immune system.^*9, 10*^ A similar strict preference for *m*-DAP containing PG fragments has also been observed for the resuscitation receptors on *B. subtilis*.^*11-13*^ Given the significant role *m*-DAP can play in microbiology and host-bacteria interactions, there is a need to synthetically build *m*-DAP based chemical probes to interrogate processes related to PG remodeling, biosynthesis, and sensing.^*14, 15*^

**Figure. 1.**
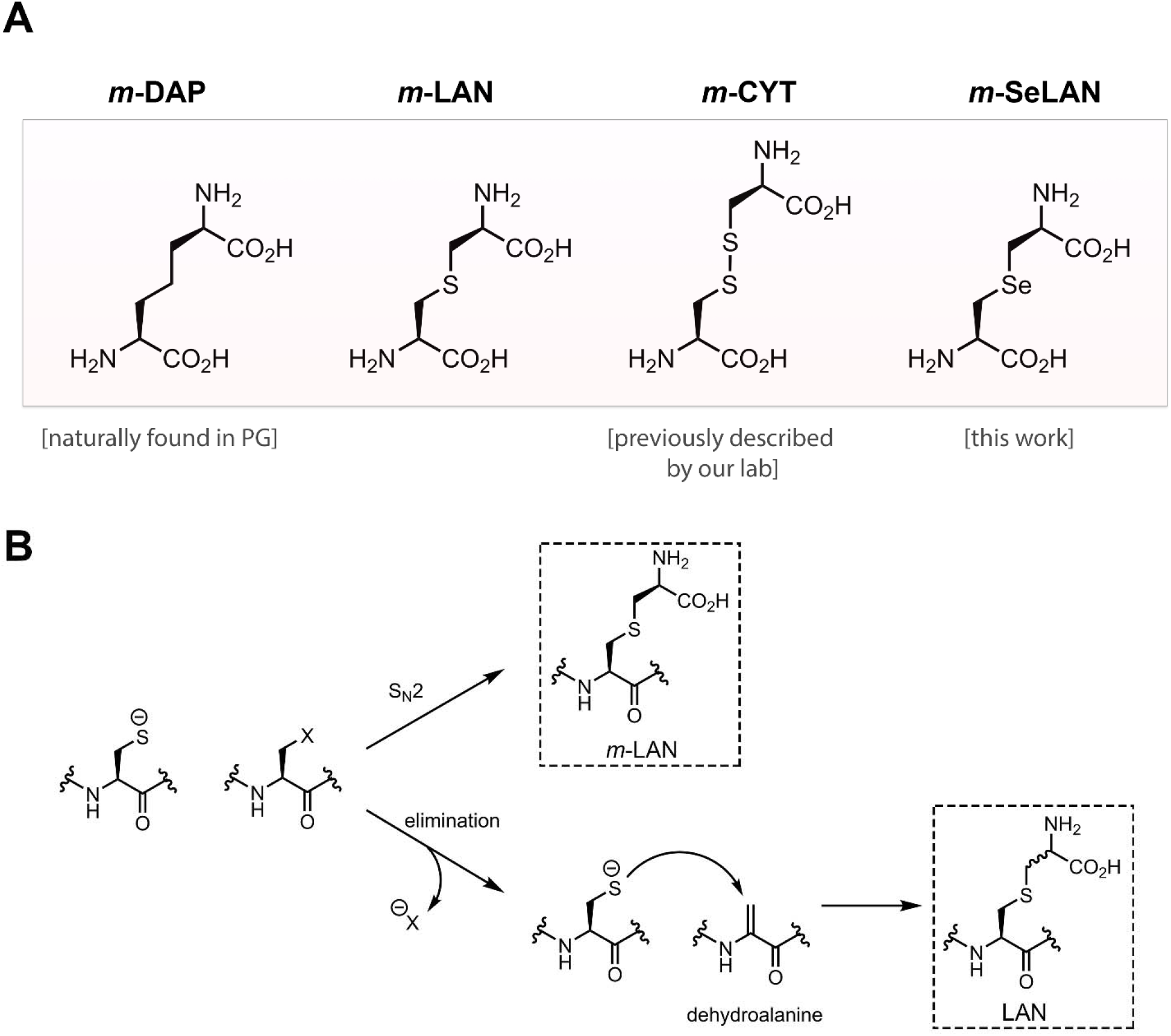
(A) Chemical structure of *m*-DAP and its three bioisosteres (*m*-LAN, *m*-CYT, and *m*-SeLAN). (B) Chemical synthesis of *m*-LAN from commercially available precursors often results in the racemization of the sidechain stereocenter due to a competing elimination reaction that occurs in the higher pH conditions required to deprotonate the sidechain of Cys.

As of today, *m*-DAP building blocks compatible with standard Fmoc-or Boc-based solid phase peptide synthesis are not commercially available. The general lack of commercial accessibility to these reagents has hampered the development of *m*-DAP based probes^*11, 16-20*^ and has, in effect, unbalanced the focus towards L-Lys containing bacterial species. A major roadblock to the generation of *m*-DAP building blocks is the presence of an internal symmetry plane, which needs to be orthogonally protected for Fmoc-based peptide synthesis.^*18*^ Due to this synthetic barrier, *m*-DAP analogues, such as *meso*-lanthionine (*m*-LAN)^*21-23*^ and *meso*-oxa-DAP, that mimic the key structural features have been described (**Fig. 1A**).^*23-26*^ Despite retaining critical structural mimicry of *m*-DAP, both *m*-LAN and *meso-*oxa-DAP have drawbacks related to complicated preparation of starting materials.

In response to this gap in readily accessible bioisosteres, we previously reported on a *m*-DAP analogue that utilized commercially available building blocks and solid-phase peptide synthesis to generate *meso*-cystine (*m*-CYT, **Fig. 1A**).^*27*^ While *m*-CYT was well tolerated as a surrogate for *m*-DAP in live bacteria, as demonstrated by its metabolic incorporation into the PG, we recognized that it may be important to retain the same number of atoms in the sidechain, which *m*-CYT does not. Moreover, the reduction of the disulfide bond in *m*-CYT may prove to be a challenge in certain conditions such as assays that require reducing agents and in intracellular compartments of mammalian cells. Motivated by the shortcomings of current bioisosteres, we sought to leverage the unique reactivity of selenium to assemble a novel *m*-DAP analog, selenolanthionine (*m*-SeLAN, **Fig. 1A**) using commercially available reagents.

## Results and Discussion

Nature has taken advantage of the sulfhydryl on the sidechain of cysteine to diversify natural products, including in the biosynthesis of lantibiotics. Lantibiotics are peptidic molecules produced by bacteria that are characterized by one or more cyclizations formed by the intramolecular nucleophilic attacks by cysteine (or methylcysteine) to form lanthionine (or methyllanthionine). While lanthionines within lantibiotics are precisely constructed by enzymes, it has proven to be considerably more challenging to control the stereochemical configuration using synthetic organic chemical methods. Prior synthetic efforts towards *m*-LAN using commercially available building blocks suffered from racemization problems ^28^, stemming from elimination and subsequent Michael addition, and formation of isomeric products that are challenging to separate (**Fig. 1B**).^*29*^ For these reasons, we envisioned that we could substitute the L-Cys precursor to *m*-CYT within the stem peptide with selenocysteine (Sec) during peptide synthesis. Following the release of the peptide from resin, the sidechain of the *m*-SeLAN can be installed by treatment with commercially available β-chloro-D-Ala. Key to the change in the strategy is the lower selenol *p*Ka value in Sec of ∼5 relative to the *p*Ka of thiol value in Cys, which is ∼8.^*30*^ The reduced *p*Ka means that the pH of the solution can be made more acidic, which will greatly suppress the elimination of β-chloro-D-Ala to dehydroalanine. ^*31*^

We reasoned that access to an *m*-DAP analog (*m*-SeLAN) could allow us to expand our toolbox of probes to investigate PG biosynthesis and remodeling. More specifically, we are interested in designing synthetic analogs that interrogate the bacterial crosslinking machinery. A prominent structural feature of the PG of all known bacteria is the crosslinking of neighboring stem peptides to afford covalent linkages that stabilize the PG layer. Disruption to PG crosslinking can be lethal to bacterial cells, as evidenced by the number of small molecule antibiotics (e.g., β-lactams, teixobactin) and immune system components (e.g., lysozyme) that target various steps in the crosslinking process.^*7, 32, 33*^ A better understanding of the key biological features of PG crosslinking could reveal under-explored modes of bactericidal agents and/or potentiating adjuvants. PG crosslinking is mediated by PG-associated transpeptidases, Penicillin Binding Proteins (PBPs) and L,D-transpeptidases (Ldts). After forming an acyl-intermediate with an active site residue to a stem peptide, transpeptidases promote the acyl-capture by the acyl-acceptor strand *via* a nucleophilic attack by the 3^rd^ position *m*-DAP residue (**Fig. 2A**). Our laboratory and others have previously demonstrated that synthetic stem peptide mimetics with fluorescent handles on the *N*-terminus can be accepted by PG transpeptidases, thereby becoming metabolically incorporated into the PG of live bacterial cells.^*34-37*^

**Figure 2.**
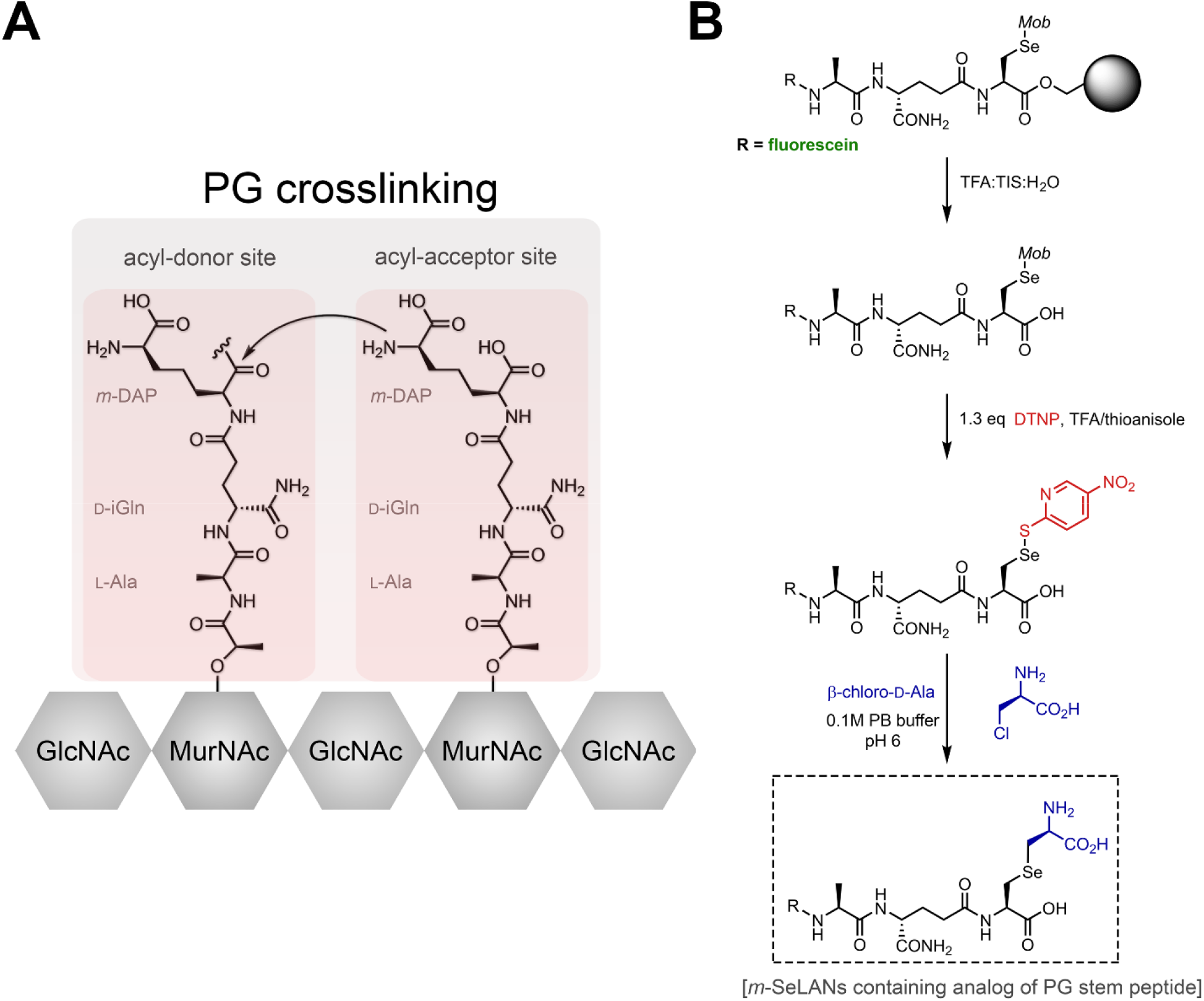
(A) In the final step of PG crosslinking by transpeptidases, an amino group from the acyl-acceptor strand performs an acyl-transfer of the acyl-donor strand. Shown is one potential configuration whereby this reaction would result in a 3-3 crosslink. Other reactions are possible based on the length of the strands in either site. (B) Synthetic route to obtain *m*-SeLAN containing peptides.

We opted to initially synthesize a tripeptide version of the stem peptide as tripeptides cannot act as acyl-donors in the PG crosslinking reactions (since they lack a terminal D-Ala necessary for the first step of the reaction) and will only participate as acyl-acceptors. Therefore, its incorporation into the PG of live bacterial cells would demonstrate the potential *m*-DAP mimicry of *m*-SeLAN. We first aimed to establish a route for the synthesis of a tripeptide mimic of the PG stem peptide using entirely commercially available reagents. To this end, the installation of Fmoc-L-Sec(Mob) at the third position could provide a site for the post-cleavage modification to provide *m*-SeLAN (**Fig. 2B**). Therefore, the amino group on the SeLAN sidechain of the tripeptide probe would be expected to act as the nucleophile in the PG crosslinking reaction. Using standard solid-phase peptide synthesis, Fmoc-L-Sec(Mob) was incorporated in third position and the *N*-terminus was modified with a fluorescein to track incorporation into the PG (fluorescence readout can be quantified by flow cytometry or confocal microscopy). After building the peptide on solid support, the Sec-containing tripeptide was released from resin using standard a standard cocktail of trifluoroacetic acid (TFA):triisopropylsilane (TIS):water (H_2_O) (9.5: 0.25: 0.25, v/v), leaving the methoxybenzyl (Mob) protecting group intact. The crude peptide was then reacted with TFA:thioanisole (9.75 : 0.25, v/v) and 1.3 equivalents of 2,2′-dithiobis(5-nitropyridine) (DTNP) to *in situ* swap out the Mob group for TNP.^*38*^ This intermediate was then allowed to react with β-chloro-D-Ala in 0.1 M phosphate buffer pH 6 and isopropanol (7:3, v/v) containing 40 equivalents of dithiothreitol (DTT).^*31*^ This reaction proceeded for 24 h at 37 ºC, was quenched with 5% TFA, and the final product (**TriSeLAN, Fig. 2B**) was purified by reversed-phase high performance liquid chromatography (RP-HPLC). In addition to **TriSeLAN**, we synthesized a peptide with a selenocystamine (**TriSeLys**) at the third position to have the ability to test the essentiality of the carboxylic acid on the side chain in live cells. The synthesis was similarly performed with the main exception being that the Sec was reacted with 2-chloro-ethylamine instead of β-chloro-D-Ala.

With the initial seleno bioisosteres probes in hand (**Fig. 3A**), we sought to evaluate the metabolic labeling of *Mycobacterium smegmatis* (*M. smegmatis*), an organism that contains *m*-DAP in its PG.^*39*^ *M. smegmatis* is also a model organism for the infectious *Mycobacterium tuberculosis*, the causative agent of Tuberculosis (TB) that is estimated to infect one third of the world’s population.^*40-42*^ For all probes synthesized, a fluorescein was included at the *N*-terminus to measure metabolic incorporation *via* flow cytometry. To test the acceptance of *m*-SeLAN, cells in culture were incubated with the various tripeptide probes to promote their metabolic incorporation into the PG scaffold during cellular growth (**Fig. 3B**). When cells were treated with **TriSeLAN**, a marked increase in cellular fluorescence of ∼ 60-fold was observed relative to untreated cells. We had previously shown that the carboxylic acid in the sidechain of the acyl-acceptor was critical for recognition and crosslinking in *M. smegmatis*.^*27*^ To confirm those results in the context of the seleno-based probes, we synthesized two additional stem peptide analogs (**TriLys** and **TriSeLys**), in which the acyl-acceptor amino group is retained. Consistently, cellular fluorescence labeling decreased considerably for both probes (**Fig. 3B**). Furthermore, similar confocal analysis with another bacterium within the same genus, *Mycobacterium bovis bacillus Calmette-Guérin* (BCG), revealed extensive labeling (**Fig. S1**). Metabolic tagging was also observed for another bacterial species whose PG scaffold contains *m*-DAP, *Lactobacillus plantarum* (**Fig. S2**). Taken together, these results are suggestive of proper mimicry of *m*-SeLAN in the context of transpeptidase processing in live cells and that the carboxylic acid is critical for metabolic labeling.

**Figure 3.**
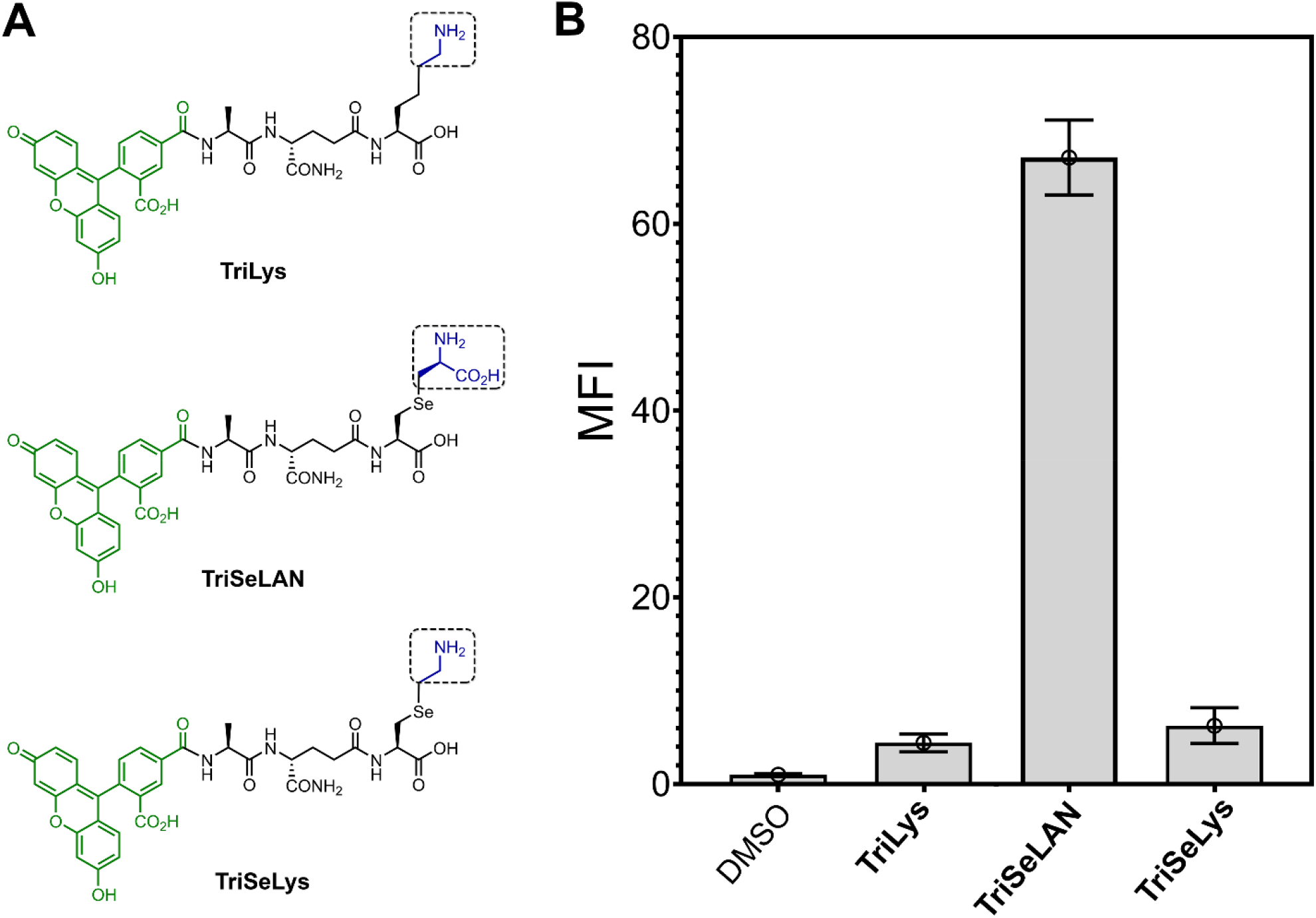
(A) Chemical structures of the tripeptides probes used in the study. The box highlights the structural differences at the crossbridge. Fluorescein is present for quantification of incorporation. (B) Flow cytometry analysis of *M. smegmatis* treated overnight with DMSO or 100 µM of designated tripeptide probes. Data are represented as mean + SD (n = 3).

Additionally, a number of experiments were performed to demonstrate the incorporation of the synthetic stem peptide analog within the PG of cells. We initially turned to SaccuFlow, a protocol we described recently to quantitatively measure the fluorescence of isolated sacculi (**Fig. 4A**).^*43*^ Sacculi from *M. smegmatis* treated with **TriSeLAN** showed high levels of fluorescence, which was consistent with the results obtained with whole cells (**Fig. 4B**). The sacculi isolation protocol is expected to remove all biomacromolecules outside of PG, which indicates that the fluorescence signal is retained even after going through the individual isolation steps. These results were confirmed by confocal imaging of the sacculi (data not shown). Metabolic incorporation into the PG scaffold was confirmed by directly analyzing muropeptides. *M. smegmatis* cultures supplemented with **TriSeLAN** were subjected to PG isolation and analyzed by liquid chromatography-mass spectrometry (LC-MS) for potential crosslinks of the fluorescent probe to endogenous PG stem peptides. We identified natural muropeptides with *N*-acetylated muramic acid as well as the *N*-glycolylated muramic sugar that *M. smegmatis* is known to present.^*44*^ The GlcNAc-MurNAc tetrapeptide (with iso-D-Glu and non-amidated *m*-DAP) as well as the N-glycoylated MurNAc tetrapeptide (with either iso-D-Glu and amidated *m*-DAP or iso-D-Gln and non-amidated *m*-DAP) were identified to be crosslinked to the fluorescent seleno-containing tripeptide, with isotopic patterns representative of selenium (**Fig. S3**).

**Figure 4.**
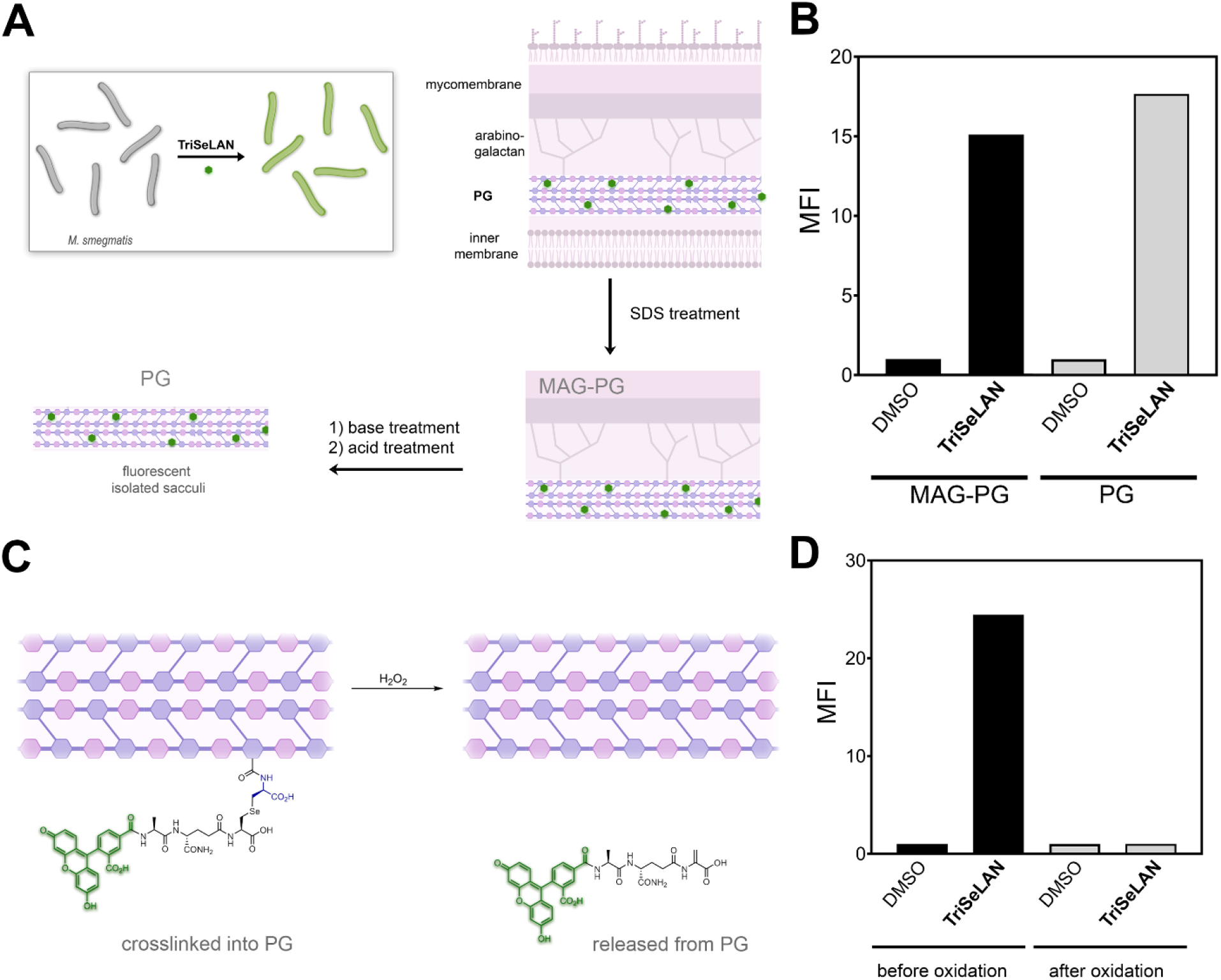
(A) treatment of *M. smegmatis* with **TriSeLAN** should result in PG tagging with a fluorescent handle. Cells are subjected to a series of PG isolation steps, including an intermediate MAG-PG and the intact final PG scaffold. (B) *M. smegmatis* treated overnight with DMSO or 100 µM of designated tripeptide probes. Flow cytometry measurements were made at intermediate points and at the final isolated PG. Data are shown for 10,000 individual sacculus. (C) Schematic diagram showing the crosslinked probe prior to and after the oxidation by hydrogen peroxide to yield a dehydroalanine (and other released products). (D) Flow cytometry analysis of metabolically tagged sacculus before and after oxidation. Data are shown for 10,000 individual sacculus.

A unique feature of selenoether (as in the one found in *m*-SeLAN) is its ability to be cleaved using orthogonal oxidation conditions to yield dehydroalanine (**Fig. 4A**).^*45, 46*^ To further confirm the presence of our probe within the PG, metabolically tagged sacculi was treated with hydrogen peroxide to induce the release of the fluorescent tag. A considerable decrease in sacculi fluorescence was measured after hydrogen peroxide treatment (**Fig. 4B**). As a control, sacculi from *M. smegmatis* treated with a non-releasable stem peptide analog did not change after exposure to the same oxidation step (**Fig. S4**). This selective oxidative cleavage in the background of all other biological functional groups has been exploited in chemoproteomics programs.^*47*^ Likewise, we envision that *m*-SeLAN may provide a handle to reverse the PG crosslinking in isolated sacculi, and potentially in live bacteria cells.

To explore the influence of *m*-SeLAN on PG crosslinking, we constructed three computational models to examine the configuration (and its impact) of natural *m*-DAP as compared to *m*-CYT and *m*-SeLAN in the active site of an L,D-transpeptidase from *M. smegmatis*. Initially, it should be pointed out that starting points for MM and QM/MM simulations were obtained from our previous study.^*27*^ In our previous QM/MM mechanistic studies involving Ldt systems,^*48*^ we described that the catalytic mechanism starts with the formation of the ionic-pair, by a proton transfer from Cys346 to His328. Therefore, we started our calculation from product state of Ldt inhibition reaction with ionic-pair of Cys/His catalytic residues. To test the influence of *m*-DAP bioisosteres, three systems were built: (I) Ldt-*m*-DAP, (II) Ldt-*m*-CYT and (III) Ldt-*m*-SeLAN (**Fig. 5A**).

**Figure 5.**
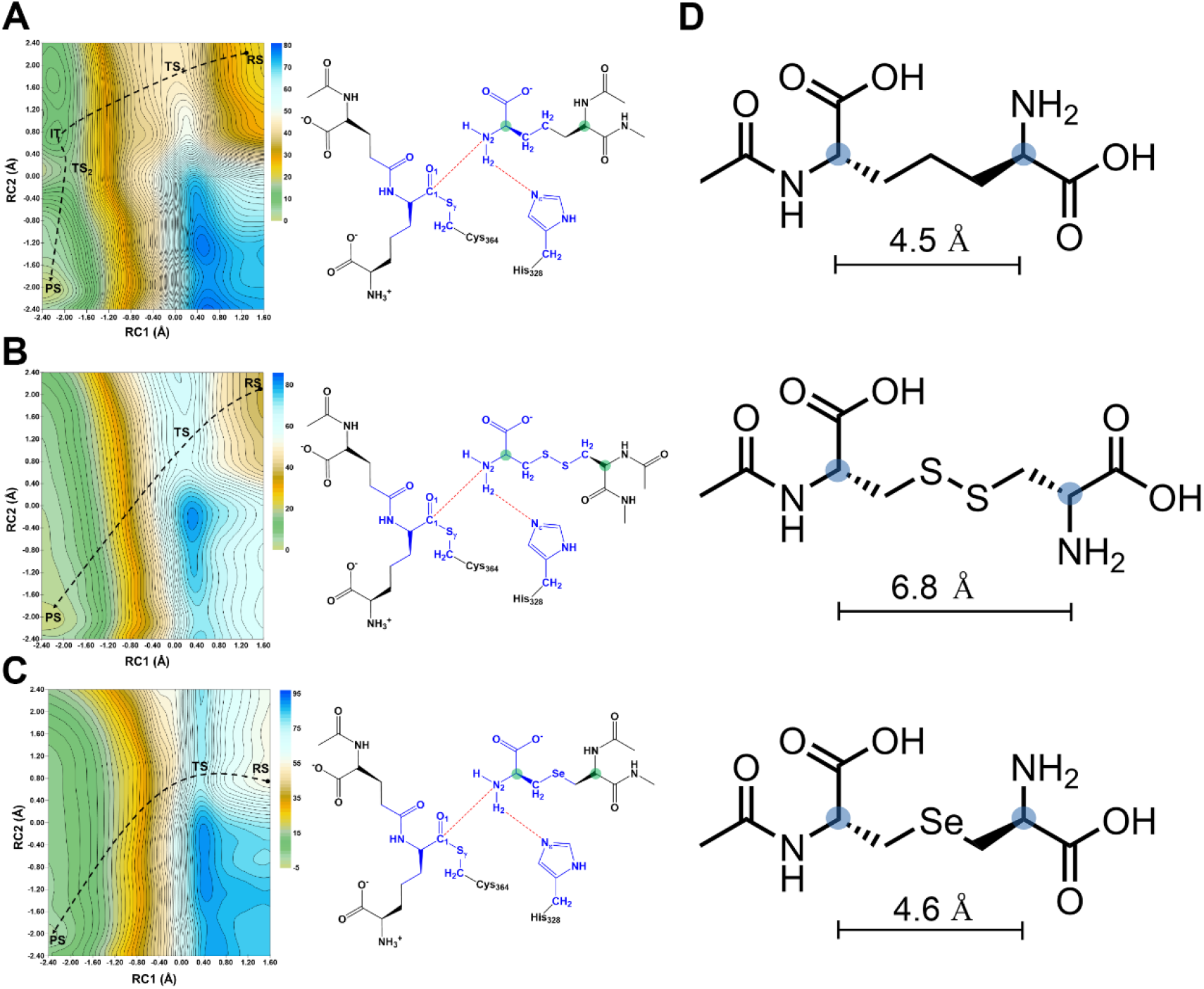
Plots of 2D-PMF at PM3/MM level (left) and combinations of RC1 and RC1 reaction coordinates (right), describing the PG cross-linking mechanism of (A) Ldt-*m*-DAP system I, (B) Ldt-*m*-CYT system II, and (C) Ldt-*m*-SeLAN system III. QM atoms for each system are in blue color. Alpha carbons of the peptide sidechain are highlighted in green color. The energy values are reported in kcal/mol. (D) Average distances from QM/MM for the designated atoms in the three bioisosteres studied.

The most relevant structural and energetic values obtained from QM/MM umbrella sampling simulations are summarized on **Table S1**. As shown in **Fig. 5A** and **Table S1**, the crosslinking mechanism of Ldt_Msm_ at PM3/MM level occurs *via* a concerted reaction for the *m*-CYT and *m*-SeLAN systems. However, for the *m*-DAP system, the reaction occurs by stepwise mechanism. In the first step, we can observe a nucleophilic attack from amine nucleophile (N_2_ atom) to carbonyl group of acyl-enzyme (C_1_ atom), which shows a theoretical free energy barrier equal to 22 kcal/mol. After, a proton transfer from amine nucleophile to imidazole group of catalytic His328 with a very small free energy barrier (about 3.4 kcal/mol). The first step of *m*-DAP system mechanism is energetically and structurally similar to reaction for the *m*-CYT and *m*-SeLAN systems.

Interestingly, from the thermodynamics perspective of the computed reaction, we observe that ΔG^R^ of *m*-SeLAN system (–55.4 kcal/mol) is much more favorable than for *m*-DAP and *m*-CYT systems (–22.7 and –37.0 kcal/mol, respectively). To explain the differences in the ΔG^R^’s, we computed the Mulliken charges (averaged over PM3/MM US frames) in the chemical species of the reactions (**Table S1**). The charges of the Sγ and C_1_ atom are similar for all systems when the RS configuration is considered. However, the charge of the N_2_ atom (amine nucleophile) has a larger electron density for *m*-SeLAN system, which allows a more favorable nucleophilic attack. At the end of reaction (PS) in all 3 systems, N_2_ atom has the same electron density. Additionally, it was observed that the distance between alpha carbons (**Fig. 5A**) are quite similar to *m*-DAP and *m*-SeLAN systems, which may explain the tolerability of PG crosslinking enzymes to this bioisostere (**Fig. 5B**).

Finally, we sought to test the tolerability of mycobacteria to the new *m*-DAP bioisostere, *m*-SeLAN, by supplementing it to cells that are auxotrophic for *m*-DAP. Uniquely, the metabolism of single amino acid m-DAP bioisosteres requires that every biosynthetic step (starting with MurE ligase) within the PG biosynthesis accommodates the synthetic analog. There is precedence in *E. coli* that are auxotrophic for *m*-DAP to have growth rescued with lanthionine (and cystathionine) supplementation.^*49, 50*^ Moreover, an unusual mutation led to a *m*-DAP auxotrophic strain of *M. smegmatis* to biosynthesize lanthionine and utilize it in place of *m*-DAP for PG biosynthesis.^*51*^ Cell growth rescue experiments were performed with *m*-DAP auxotrophic *M. smegmatis* cells (**Fig. 6**) by adopting prior protocols. Briefly, cells were grown to early log (OD_600_ ∼0.2), then the media was supplemented with *m*-DAP alone or a combination of *m*-DAP and *m*-SeLAN. In the absence of supplementation of *m*-DAP, the optical density does not change over 24 h post media exchange. As expected, addition of *m*-DAP to the media rescues growth at both 5 and 10 µg/mL. Supplementation of *m*-SeLAN led a measurable increase in optical density when co-incubated with 5 µg/mL of *m*-DAP, which indicates the proper utilization of this bioisostere in *M. smegmatis m*-DAP auxotrophs.

**Figure 6.**
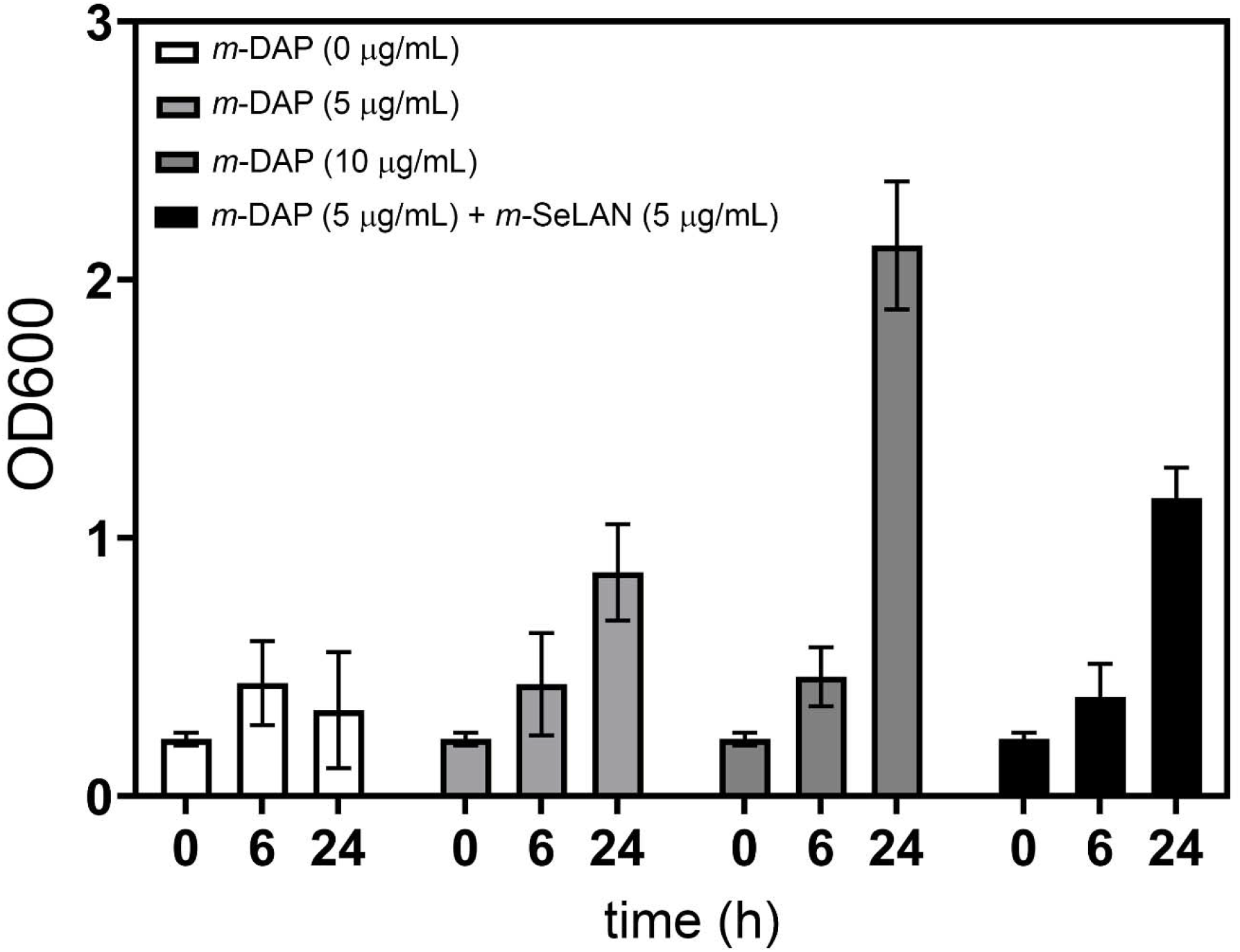
*m*-DAP auxotrophic *M. smegmatis* were grown to OD_600_ of 0.25 in media supplemented with exogenous *m*-DAP, at which point the media was exchanged with the media described for each condition. The OD_600_ was then measured at time 0, 6, and 24 h post media exchange. Data are represented as mean + SD (n = 3).

## Conclusion

Here, we described a strategy that uses solid-phase peptide synthesis to construct a selenolanthionine *m*-DAP mimic. This probe was tolerated by endogenous PG crosslinking machinery in live *M. smegmatis* and BCG cells, as investigated by the use of flow cytometry and confocal microscopy. Importantly, the *m*-SeLAN tripeptide probe specifically suggests proper *m*-DAP mimicry, as the tripeptide probe acts as an acyl-acceptor in the PG crosslinking step. The incorporation of selenolanthionine based probes was also observed in other organisms that incorporate *m*-DAP within their PG. A number of lines of evidence pointed to the incorporation of this probe within the PG of mycobacteria, including an assay in which the selenolanthionine was selectively released from the isolated PG. *m*-SeLAN was further shown to have *m*-DAP mimicry when *m*-DAP auxotrophs were rescued by supplementation of single amino acid *m*-SeLAN and *m*-DAP into the media. We envision that these probes can provide an alternate synthetic route for *m*-DAP bioisosteres that involves a selenoether linkage (established by the use of completely commercial reagents) which prevents the possibility of reduction and maintains the approximate length of the *m*-DAP sidechain, expanding the tool set currently being used to study PG biosynthesis.

## Supporting Information

Additional figures, tables, and materials/methods are included in the supporting information file.

