## Supporting Information for "Metabolic Processing of Selenium-based Bioisostere of *meso*-diaminopimelic Acid in Live Bacteria"

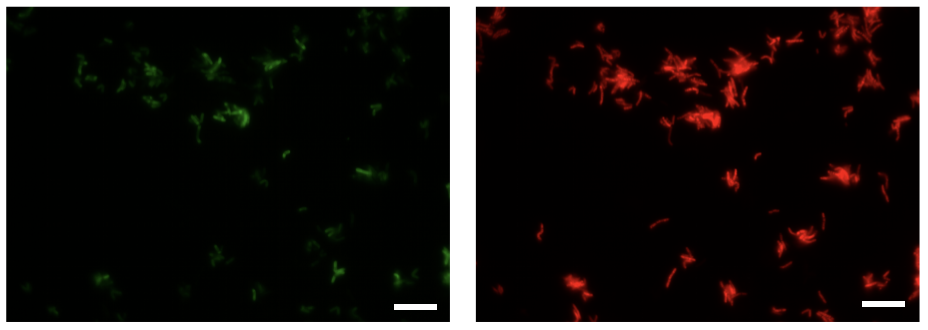

**Figure S1.** Confocal microscopy image of BCG cells treated with **TriSeLAN** (200 μM). Cells express mCherry. Green fluorescence images (left) show that the probe is associated with cells and they co-localize with the mCherry signal. Scale bar = 10 μm.

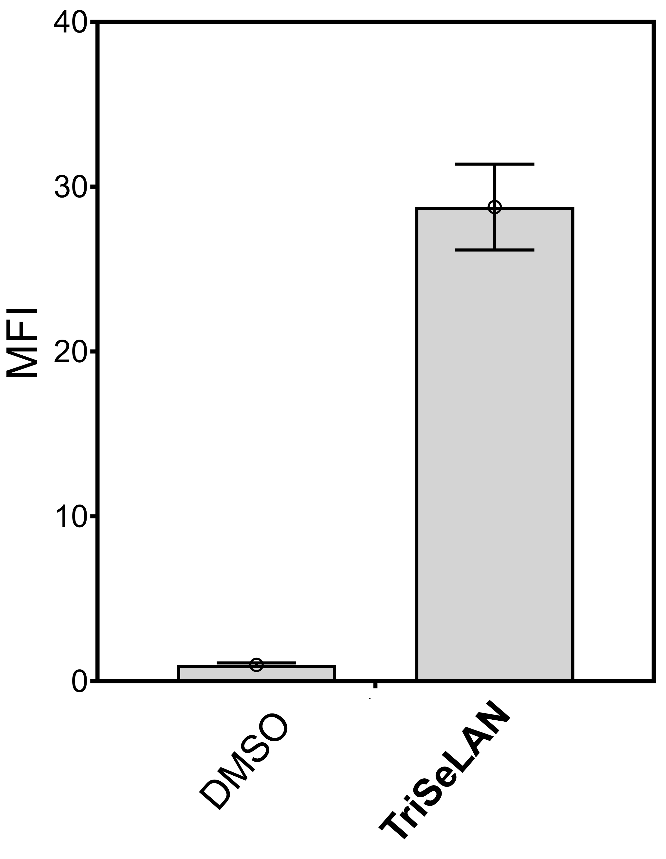

**Figure S2.** Flow cytometry analysis of *L. plantarum* treated overnight with DMSO or 100 µM of designated tripeptide probes. Data are represented as mean + SD (n = 3).

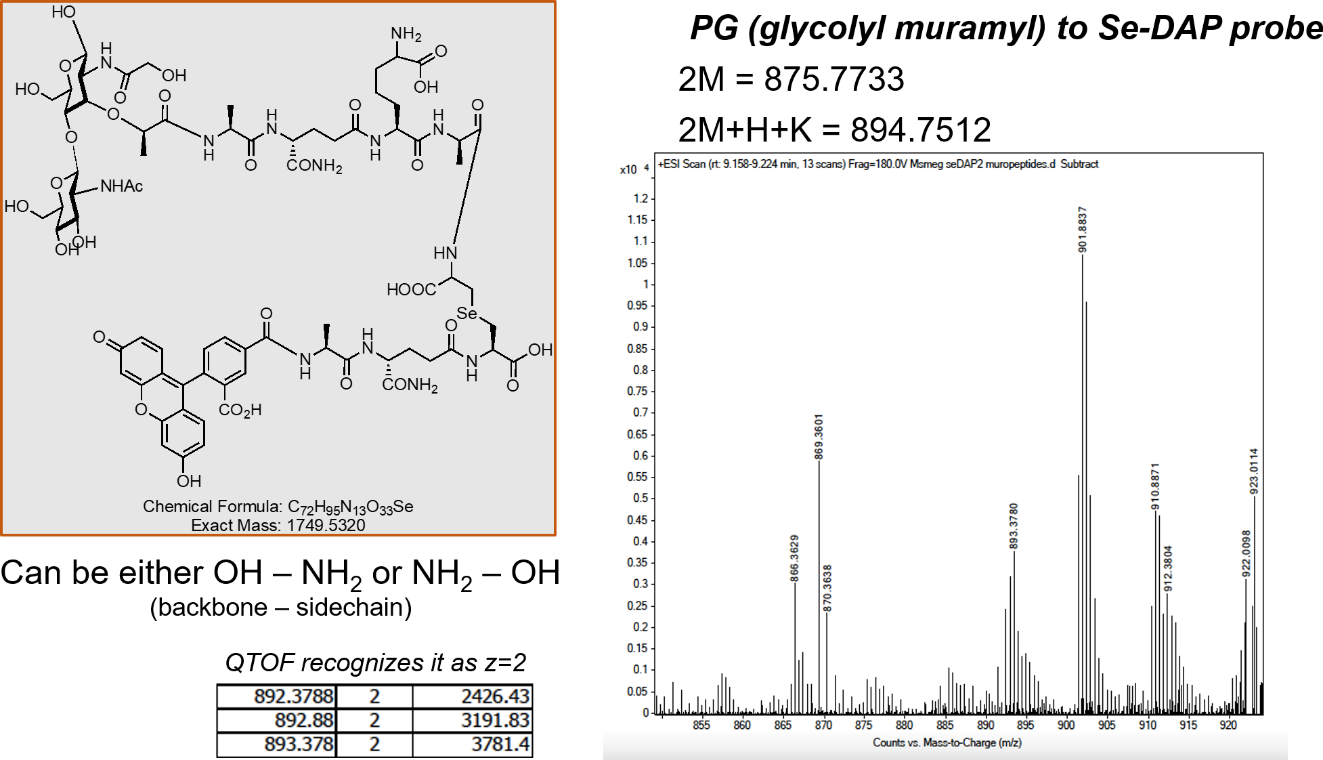

**Figure S4.** Mass spectrometry analysis of fragments of PG isolated from M. smegmatis treated overnight with **TriSeLAN**.

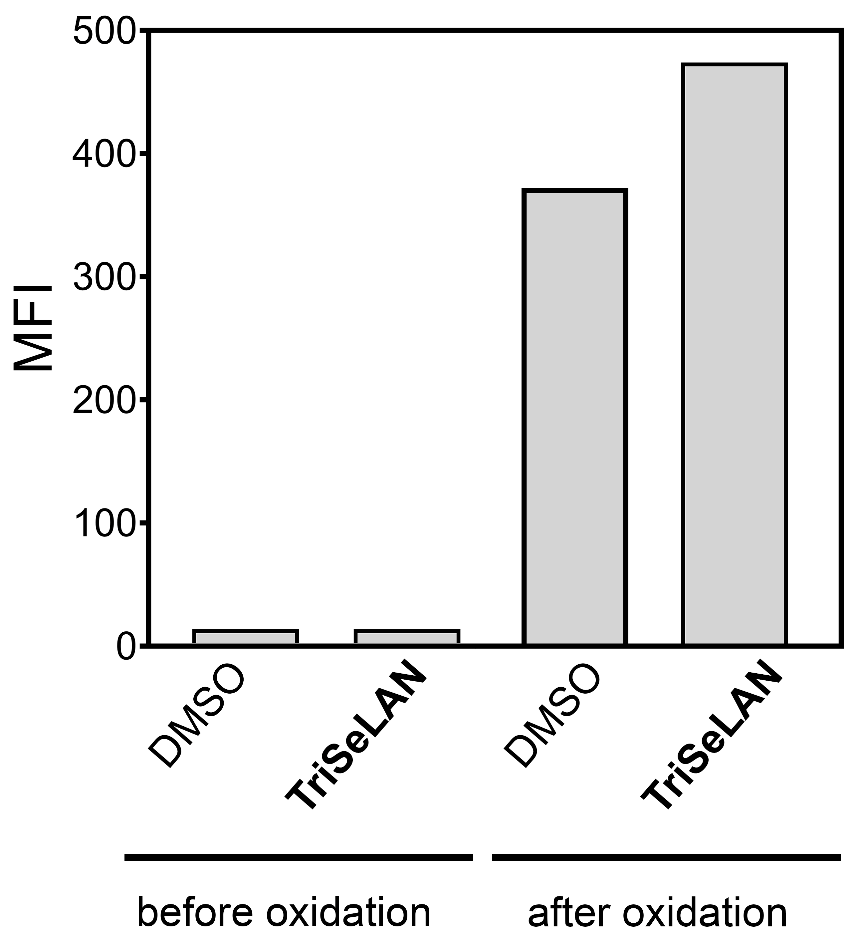

**Figure S4** Flow cytometry analysis of sacculi isolated from *M. smegmatis* treated overnight with DMSO or 100 µM of **Fl-TriLys**. Data are represented as mean + SD (n = 3).

**Table S1.** Averaged of Key Distances (in Å) and Thermodynamics Parameters (in kcal/mol) obtained by PM3/MM 2D-PMF. The standard deviation values are in parenthesis. The 3D structure (PDB format) of each PS is available on SI file.

| **m-DAP system** | | | | | | | |
| --- | --- | --- | --- | --- | --- | --- | --- |
| distance | RS | TS_1_ | IT | TS_2_ | PS | ΔG^‡^ | ΔG^R^ |
| d(S_γ_–C_1_) | 1.83  (0.04) | 2.10  (0.04) | 3.72  (0.05) | 3.72  (0.05) | 3.72  (0.05) | 22.0/3.4 | –22.7 |
| d(N_2_–C_1_) | 3.44  (0.05) | 1.94  (0.05) | 1.53  (0.04) | 1.41  (0.03) | 1.41  (0.03) |  |  |
| d(N_2_–H_2_) | 1.00  (0.03) | 1.02  (0.03) | 1.06  (0.03) | 1.35  (0.05) | 3.03  (0.04) |  |  |
| d(N_ε_–H_2_) | 3.20  (0.04) | 3.00  (0.04) | 1.85  (0.04) | 1.42  (0.04) | 1.02  (0.03) |  |  |
| d(Cα) | 4.80  (0.24) | 4.50  (0.14) | 5.03  (0.13) | 5.10  (0.07) | 4.96  (0.10) |  |  |
| **m-CYT system** | | | | | | | |
| distance | RS | TS | PS |  |  | ΔG^‡^ | ΔG^R^ |
| d(S_γ_–C_1_) | 1.88  (0.04) | 2.15  (0.05) | 3.62  (0.04) |  |  | 26.4 | –37.0 |
| d(N_2_–C_1_) | 3.50  (0.05) | 2.22  (0.06) | 1.42  (0.04) |  |  |  |  |
| d(N_2_–H_2_) | 0.99  (0.03) | 1.35  (0.03) | 3.77  (0.12) |  |  |  |  |
| d(N_ε_–H_2_) | 2.99  (0.05) | 2.23  (0.04) | 1.08  (0.11) |  |  |  |  |
| d(Cα) | 7.33  (0.12) | 6.83  (0.10) | 6.62  (0.27) |  |  |  |  |
| **SeLAN system** | | | | | | | |
| distance | RS | TS | PS |  |  | ΔG^‡^ | ΔG^R^ |
| d(S_γ_–C_1_) | 1.89  (0.04) | 1.96  (0.05) | 3.83  (0.04) |  |  | 20.2 | –55.4 |
| d(N_2_–C_1_) | 3.29  (0.05) | 2.40  (0.05) | 1.42  (0.03) |  |  |  |  |
| d(N_2_–H_2_) | 1.04  (0.02) | 1.54  (0.03) | 3.75  (0.09) |  |  |  |  |
| d(N_ε_–H_2_) | 2.80  (0.03) | 1.84  (0.04) | 1.00  (0.04) |  |  |  |  |
| d(Cα) | 4.28  (0.15) | 4.60  (0.13) | 4.88  (0.12) |  |  |  |  |

| **m-DAP system** | | | | | |
| --- | --- | --- | --- | --- | --- |
| Atom | RS | TS_1_ | IT | TS_2_ | PS |
| S_γ_ | –0.001  (0.004) | –0.234  (0.005) | –0.982 (0.003) | –0.112 (0.004) | –0.969 (0.002) |
| C_1_ | 0.221  (0.003) | 0.314  (0.003) | 0.213  (0.003) | 0.265  (0.002) | 0.265 (0.002) |
| N_2_ | –0.007  (0.002) | 0.141  (0.005) | 0.417  (0.005) | –0.002  (0.003) | –0.002  (0.004) |
| **m-CYT system** | | | | | |
| Atom | RS | TS | PS |  |  |
| S_γ_ | –0.005  (0.004) | –0.143  (0.004) | –0.330  (0.006) |  |  |
| C_1_ | 0.211  (0.003) | 0.339  (0.002) | 0.263  (0.003) |  |  |
| N_2_ | 0.329  (0.006) | –0.007  (0.003) | –0.002  (0.004) |  |  |
| **SeLAN system** | | | | | |
| Atom | RS | TS | PS |  |  |
| S_γ_ | –0.005  (0.004) | –0.140  (0.005) | –0.261  (0.006) |  |  |
| C_1_ | 0.221  (0.003) | 0.299  (0.002) | 0.248  (0.002) |  |  |
| N_2_ | –0.125  (0.002) | –0.134  (0.002) | –0.003  (0.003) |  |  |

**Materials.** All peptide related reagents (resin, coupling reagent, deprotection reagent, amino acids, and cleavage reagents) were purchased from ChemImpex. Bacterial strains *M. smegmatis* WT (14468) were grown in Middlebrook 7H9 media supplemented with 0.05% Tween 80 and enriched with albumin/dextrose/catalase (ADC) and glycerol (0.2% v/v).

**Flow cytometry analysis of bacterial labeling.** Media containing 100 µM of each probe ­**­­­­­­**­­were prepared. Bacterial cells from an overnight culture were added to the medium (1:100 dilution) and allowed to grow overnight at 37°C with shaking at 250 rpm. The bacteria were harvested at 6,000g and washed three times with original culture volume of 1X PBS followed by fixation with 2% formaldehyde in 1X PBS for 30 min at room temperature. The cells were washed once more to remove formaldehyde and then analyzed using an Attune NxT flow cytometer equipped with a 488 nm laser and 525/40 nm bandpass filter. The data were analyzed using the Attune NxT Software, where populations were gated and no less than 10,000 events per sample were recorded.

**Muropeptide Isolation of *M. smegmatis*.** M7H9 media ADC enriched containing 0.05% Tween 80 and 0.2% glycerol was aliquoted into a 96-well plate as either blank (no probe) or with 100 µM of fluorescent probe (Se-containing tripeptide) following a protocol established in the literature to miniaturize Mycobacterium muropeptide isolation.[1, 2] *M. smegmatis* bacteria were added to the medium (1:100) and allowed to grow overnight at 37°C with shaking at 250 rpm. Cells were then harvested at 2,700g for 15 min at 4°C and washed 1X in a minimal volume of 1X PBS. Pellets were resuspended in 10 mM NH_4_HCO_3_ with a protease inhibitor cocktail (Sigma SRE005- 1BO). This suspension was lysed using a tip sonicator (Fisher Scientific) at 60% amplitude for 60 seconds, with at least 60 seconds on ice in between. This was repeated for 5 cycles. The sonicate was digested with 10 µg/mL DNAse and RNase for 1 h at 4°C, then harvested at 2,700g for 30 min at 4°C. The pellet was resuspended in PBS with 2% SDS and incubated for 1 h at 50°C with constant stirring. This was collected using the same centrifugal parameters and the SDS process was repeated 2X. The resulting pellet was resuspended in PBS with 1% SDS and 0.1 mg/mL Proteinase K at 45°C for 1 h with stirring. The sample was then heated to 90°C for 1 h and collected as above. This step at 90°C was repeated 2X. The sample was washed 2X with PBS and 4X with dH2O. This yielded mycolic-arabinogalactan-peptidoglycan (mAGP). To remove the mycolic acid layer, samples were collected and resuspended in 0.5% KOH in MeOH and incubated at 37°C for 4 days with shaking. After 4 days, samples were collected, washed twice with MeOH and twice with diethyl ether to yield the arabinogalactan layer. To remove the arabinogalactan layer, samples were resuspended in 0.5 N H_2_SO_4_ at 37°C for 5 days with shaking. After 5 days, samples were collected, washed 4X with dH_2_O and lyophilized. The lyophilized sacculi was resuspended in 0.5 mL of 10 mM sodium acetate pH 5 with 25 µg of muramidase and incubated at 37°C for 16 h with shaking. Sacculi was spun down, the supernatant was filtered (10 kDa) cutoff and dried under vacuum. Dried sacculi was dissolved in HPLC-grade water and an aliquot was applied to a C18(2) column (Luna 5 µm 100Å 250 x 4.6 mm) connected to an Agilent LC-QTOF (Agilent 1260 Infinity II Prime LC with Agilent 6545B QTOF). Muropeptide samples were eluted with a 2 to 30% linear gradient of water to acetonitrile (0.5% formic acid) at 0.4 mL/min. The muropeptides were analyzed by MS using MassHunter software.

**BCG Confocal Imaging.** For imaging BCG: mCherry-expressing *Mycobacterium bovis* BCG Tokyo was grown in 7H9 medium at 37 C to an OD600 of 0.3 then incubated with 200 µM  FL-**TriQSeLAN** for an additional 24 hrs at 37 C. Bacteria were washed 3x in PBS + 0.01% bovine serum albumin + 0.05% Tween-80 (PBSTB) then fixed with 4% formaldehyde for 2 hrs at room temperature. Samples were washed an additional 3x with PBSTB and stored at 4 C prior to imaging.

**m-DAP auxotrophic growth analysis.** A diaminopimelate auxotroph (strain mc21620) was grown at 30 C in 7H9 medium supplemented with l-lysine, l-methionine, and l-threonine at 40 mg/ml and dl-homoserine at 80 µg/ml as previously described (see reference below). A primary culture was grown in the presence of 200 µg/ml DAP, then washed and resuspended at an OD600 of 0.2 in the supplemented 7H9 medium +/- DAP +/- the compounds synthesized in this manuscript at the indicated concentrations.

**Scheme S1. Synthesis of TriSe(Mob).**

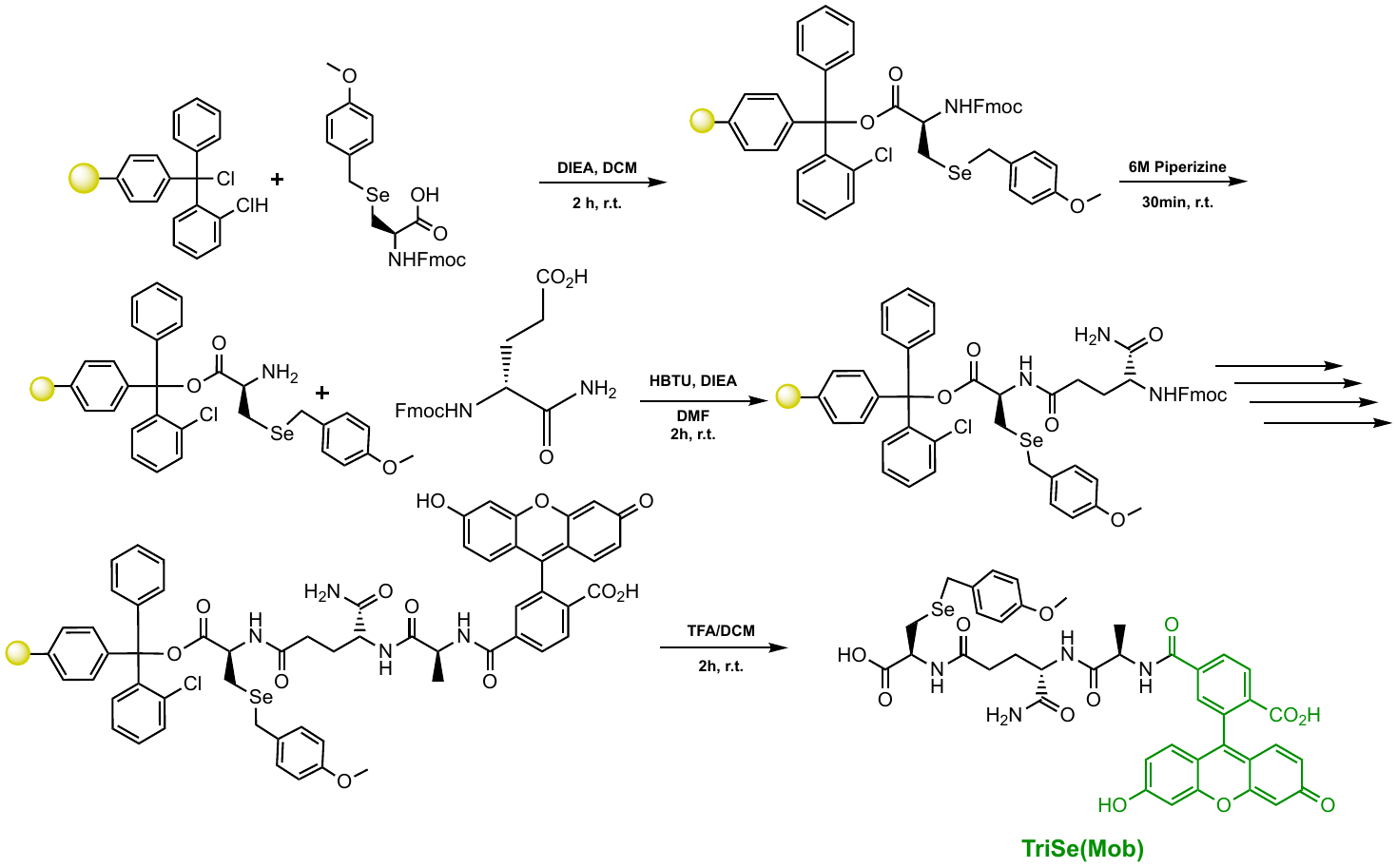

Fmoc-L-Sec(Mob) (1.1 eq, 154 mg, 0.303 mmol) was added to a 25 mL peptide synthesis vessel charged with 2-chlorotrityl chloride resin (250 mg, 0.275 mmol) and DIEA (4 eq, 0.191 mL, 1.10 mmol) in dry DCM (5 mL). The resin was agitated for 2 h at ambient temperature and washed with MeOH and DCM (3 x 15 mL each). The Fmoc protecting group was removed with 6 M piperazine/100 mM HOBt in DMF (15 ml) for 30 min at ambient temperature, then washed as before. Fmoc-D-glutamic acid α-amide (3 eq, 304 mg, 0.825 mmol), HBTU (3 eq, 314 mg, 0.825 mmol), and DIEA (6 eq, 0.287 mL, 1.65 mmol) in DMF (10 mL) were added to the reaction flask and agitated for 2 h at ambient temperature. The Fmoc deprotection and coupling procedure was repeated as before using the same equivalencies with Fmoc-L-Alanine-OH. The Fmoc group of L-alanine was deprotected and coupled with 5(6)-carboxyfluorescein (2 eq, 207 mg, 0.55 mmol), HBTU (2 eq, 209 mg, 0.55 mmol) and DIEA (4 eq, 0.191 mL, 1.10 mmol) in DMF (10 mL) shaking overnight. The resin was washed as before and added to a solution of TFA/H_2_O/TIPS (95%, 2.5%, 2.5%, 20 mL) with agitation for 2 h at ambient temperature. The resin was filtered and the resulting solution was concentrated *in vacuo* and trituated with cold diethyl ether, leaving the Mob protecting group intact.

**R1. Reaction to obtain TriSeLAN.**

The crude peptide from Scheme S1 (**TriSe(Mob)**) was treated with TFA:thioanisole (9.75 : 0.25, v/v) and 1.3 equivalents of 2,2′-dithiobis(5-nitropyridine) (DTNP) for 1 h at room temperature to *in situ* swap out the Mob group for TNP. This intermediated was then allowed to react with β-chloro-D-Ala in 0.1 M phosphate buffer pH 6 and isopropanol (7:3, v/v) containing 40 equivalents of dithiothreitol (DTT). This reaction proceeded for 24 h at 37 ºC, was quenched with 5% TFA, and isopropanol was removed *in vacuo*. The final product was then purified using RP-HPLC with H_2_O/MeOH and analyzed for purity using a Waters 1525 pump and 2489 detector equipped with a Phenomenex Luna 5µ C8(2) 100Å (250 x 4.60 mm) column; gradient elution with H_2_O/CH_3_CN.

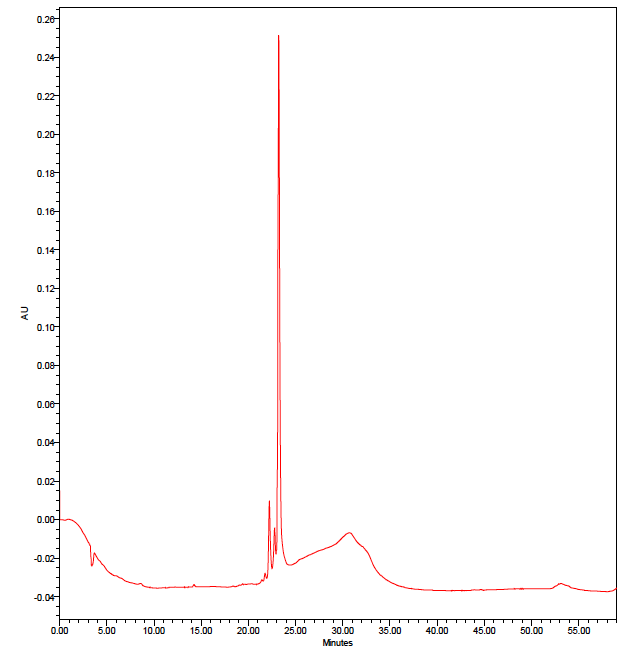

ESI-MS calculated [M + H^+^]: 814.1470, found: 814.1362

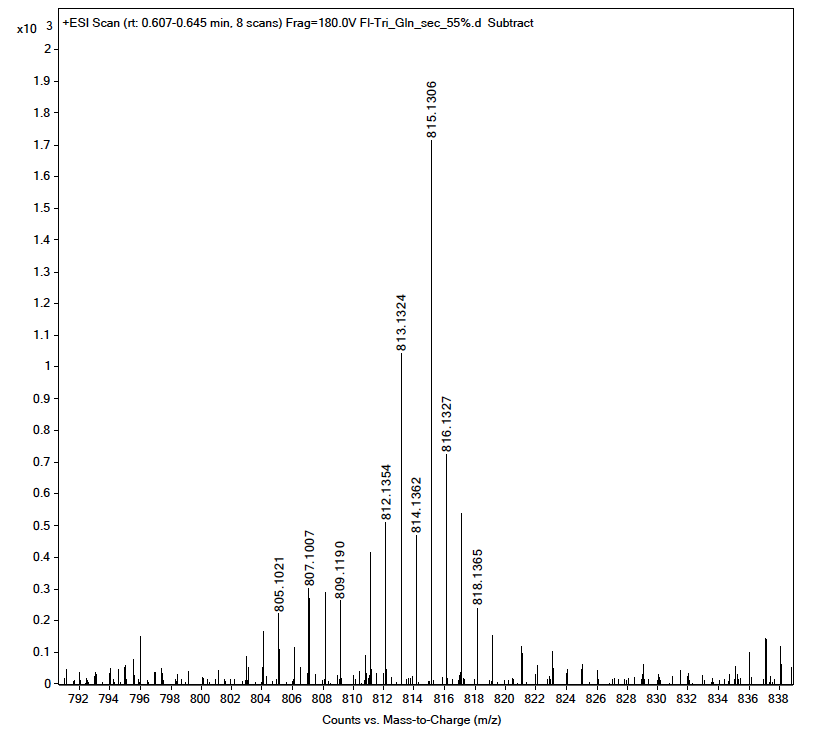

**R2. Reaction to obtain TriSeLys.**

The crude peptide from Scheme S1 (**TriSe(Mob)**) was treated with TFA:thioanisole (9.75 : 0.25, v/v) and 1.3 equivalents of 2,2′-dithiobis(5-nitropyridine) (DTNP) for 1 h at room temperature to *in situ* swap out the Mob group for TNP. This intermediated was then allowed to react with 2-chloro-ethylamine in 0.1 M phosphate buffer pH 6 and isopropanol (7:3, v/v) containing 40 equivalents of dithiothreitol (DTT). This reaction proceeded for 24 h at 37 ºC, was quenched with 5% TFA, and isopropanol was removed *in vacuo*. The final product was then purified using RP-HPLC with H_2_O/MeOH and analyzed for purity using a Waters 1525 pump and 2489 detector equipped with a Phenomenex Luna 5µ C8(2) 100Å (250 x 4.60 mm) column; gradient elution with H_2_O/CH_3_CN.

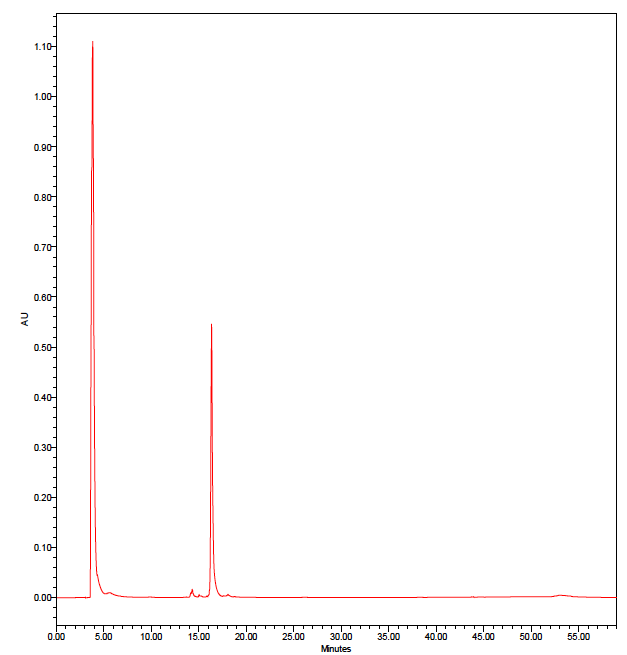

ESI-MS calculated [M + H^+^]: 770.1571, found: 770.1561

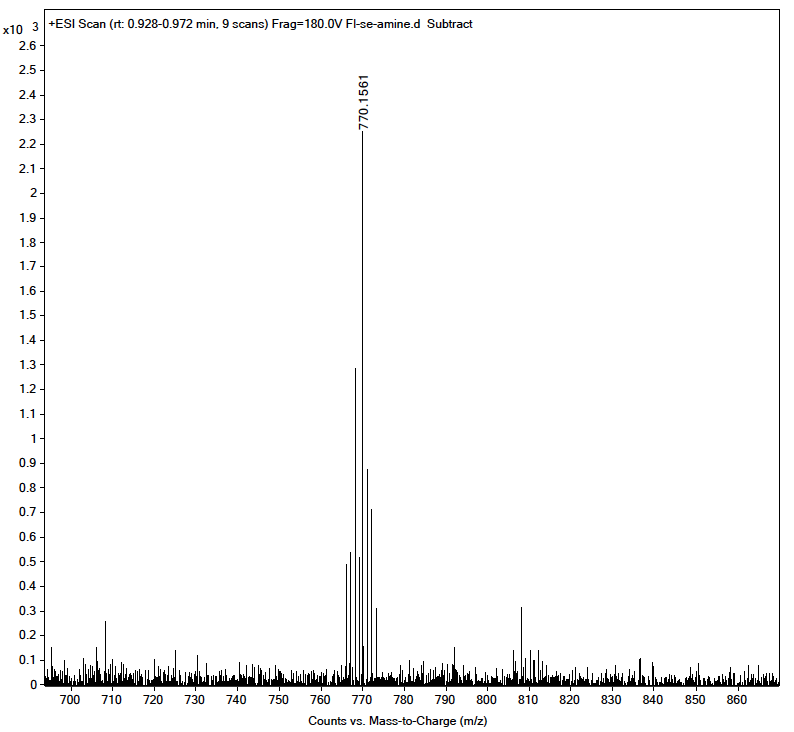

**R3. Reaction to obtain Single AA SeLAN.**

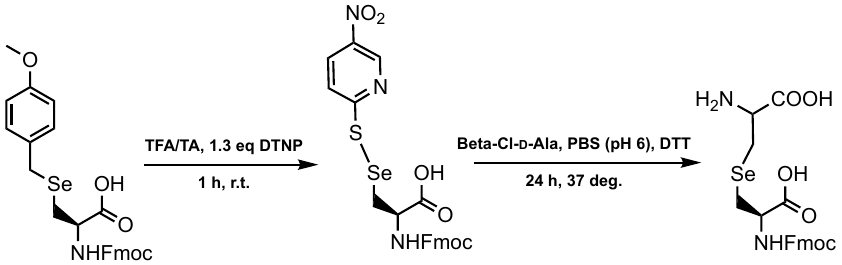

Fmoc-L-Sec(Mob) (50 mg) was treated with TFA:thioanisole (9.75 : 0.25, v/v) and 1.3 equivalents of 2,2′-dithiobis(5-nitropyridine) (DTNP) for 1 h at room temperature to *in situ* swap out the Mob group for TNP. The resulting solution was concentrated *in vacuo*, dissolved in H_2_O and lyophilized. Resulting solid Fmoc-L-Sec-TNP was reacted with β-chloro-D-Ala in 0.1 M phosphate buffer pH 6 and isopropanol (7:3, v/v) containing 40 equivalents of dithiothreitol (DTT). This reaction proceeded for 24 h at 37 ºC, was quenched with 5% TFA, and isopropanol was removed *in vacuo*. The final product (Fmoc-SeLAN) was then purified using RP-HPLC with H_2_O/MeOH. Piperazine (100 mg, 1.2 mmol) was coupled to 2-chlorotrityl chloride resin (150 mg, 0.165 mmol) for 1 h at ambient temperature with anhydrous DCM (5 mL) and DIEA (4 eq, 0.574 mL, 3.3 mmol). The resin was washed with 1% DIEA in MeOH (50 mL) and transferred to a round bottom flask with a stir bar, where the concentrated HPLC fractions were added and stirred for 30 m at room temperature to remove Fmoc. The resin was filtered and the resulting liquid was concentrated *in vacuo*.

ESI-MS calculated [M + H^+^]: 257.0035, found: 257.0022

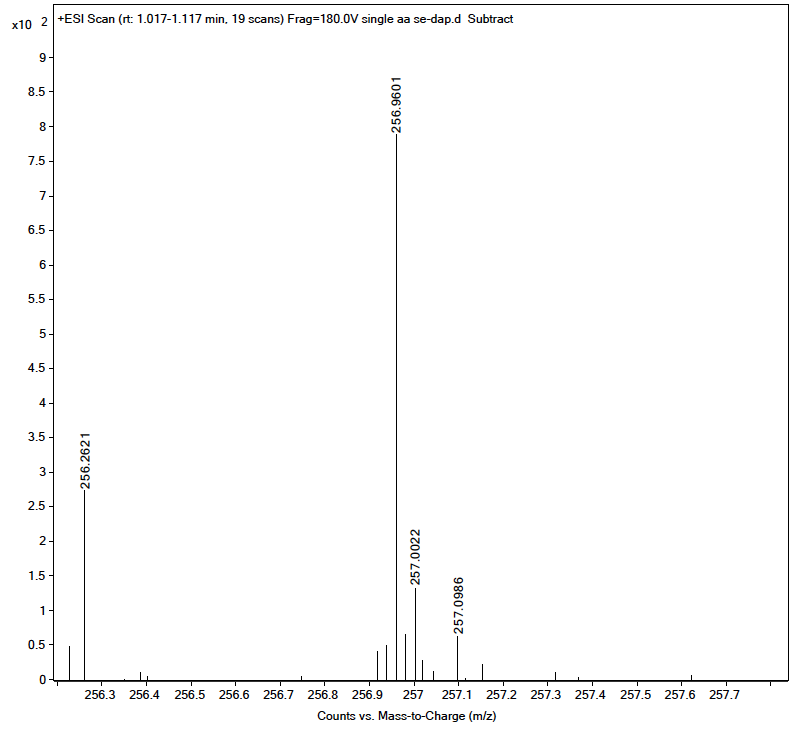

1.	Kühner, D., et al., *From cells to muropeptide structures in 24 h: Peptidoglycan mapping by UPLC-MS.* Scientific Reports, 2014. **4**(1): p. 7494.

2.	Mahapatra, S., et al., *Unique Structural Features of the Peptidoglycan of Mycobacterium leprae.* Journal of Bacteriology, 2008. **190**(2): p. 655-661.

**Computational Section**

**System setup**

The starting structure for the construction of our *M. smegmatis* LDT (LDT_Msm_) and PG fragments were obtained and *in silico* modified from our previous publication (ACS Chem. Biol. 2020, 15, 2966−2975). Particularly, SeLAN fragment was optimized at HF/6-31G* QM level by Gaussian09 package (Frisch et al. 2009) and the RESP approach (Woods and Chappelle 2000) was used to obtain the MM charges. The GAFF parameters (Journal of Computational Chemistry 25 (9): 1157–74) for SeLAN were adapted from (J. Phys. Chem. A 2016, 120, 4389−4400). Then, all LDT systems were submitted to the same classical procedure for MD simulations as described by our previous study (ACS Chem. Biol. 2020, 15, 2966−2975).

**QM/MM umbrella sampling and Potential of Mean Force (PMF)**

QM/MM MD simulations were performed out using SANDER module of AmberTools21 package (Seabra et al. 2007), where a stable representative conformation from the previous MM MD simulations was chosen QM/MM MD calculations. Here, in order to avail all system under same QM Hamiltonian, the PM3 semi-empirical method (J Comput Chem. 1989, 10, 209–264) was selected for the QM part, while the MM part was described by ff14SB (Maier et al. 2015). To satisfy the frontier on the QM and MM region, we selected the hydrogen link atoms (Field, Bash, and Karplus 1990). The QM part has a total of 66, 65 and 64 atoms for the m-DAP, m-CYT and SeLAN systems, respectively.

QM/MM umbrella sampling MD simulations were performed to obtain the 2D Potential of Mean Force (2D-PMF) of the acyl-intermediate of the LDT reaction along two reaction coordinates: RC1 = d(S_γ_–C_1_) – d(N_2_–C_1_) (from –2.40 to 1.60 Å to describe the nucleophilic attack from amine nucleophile to carbonyl group of acyl-enzyme) and RC2 = d(N_2_–H_2_) – d(N_ε_–H_2_) (from –2.40 to 2.40 Å to describe the proton transfer from amine nucleophile to imidazole group of catalytic His328) (see Figure XXX for details), each RCs were sampled in steps of 0.20 Å and their values were saved every 2 fs of interval.

Following our previous procedures, (ACS Chem. Biol. 2020, 15, 2966−2975), for each window, 2 ps of equilibration were performed before run 20 ps of production with a time step of 1 fs. Besides, during QM/MM umbrella sampling calculations, RCs were constrained by applying a harmonic potential of force constant of 250 kcal/mol∙Å^2^. Finally, the 2D-PMF of each LDT system was computed by applying the variational free energy profile (VFEP) (Lee et al. 2013) as implemented in the VFEP package developed by Lee and co-workers (Lee et al. 2014), which has been has been successfully applied for LDT systems (J.R.A. Silva, Roitberg, and Alves 2014; José Rogério A Silva et al. 2015) (ACS Chem. Biol. 2020, 15, 2966−2975).

Field, Martin J, Paul A Bash, and Martin Karplus. 1990. “A Combined Quantum Mechanical and Molecular Mechanical Potential for Molecular Dynamics Simulations.” *Journal of Computational Chemistry* 11 (6): 700–733. https://doi.org/10.1002/jcc.540110605.

Frisch, M J, G W Trucks, H B Schlegel, G E Scuseria, M A Robb, J R Cheeseman, G Scalmani, et al. 2009. “Gaussian 09 Revision A.2.”

Lee, Tai-Sung, Brian K Radak, Ming Huang, Kin-Yiu Wong, and Darrin M York. 2014. “Roadmaps through Free Energy Landscapes Calculated Using the Multidimensional VFEP Approach.” *Journal of Chemical Theory and Computation* 10 (1): 24–34. https://doi.org/10.1021/ct400691f.

Lee, Tai-Sung, Brian K Radak, Anna Pabis, and Darrin M York. 2013. “A New Maximum Likelihood Approach for Free Energy Profile Construction from Molecular Simulations.” *Journal of Chemical Theory and Computation* 9 (1): 153–64. https://doi.org/10.1021/ct300703z.

Maier, James A, Carmenza Martinez, Koushik Kasavajhala, Lauren Wickstrom, Kevin E Hauser, and Carlos Simmerling. 2015. “Ff14SB: Improving the Accuracy of Protein Side Chain and Backbone Parameters from Ff99SB.” *Journal of Chemical Theory and Computation* 11 (8): 3696–3713. https://doi.org/10.1021/acs.jctc.5b00255.

Seabra, Gustavo de M, Ross C Walker, Marcus Elstner, David A Case, and Adrian E Roitberg. 2007. “Implementation of the SCC-DFTB Method for Hybrid QM/MM Simulations within the Amber Molecular Dynamics Package.” *The Journal of Physical Chemistry A* 111 (26): 5655–64. <https://doi.org/10.1021/jp070071l>.

Silva, J.R.A., A.E. Roitberg, and C.N. Alves. 2014. “Catalytic Mechanism of L,D-Transpeptidase 2 from Mycobacterium Tuberculosis Described by a Computational Approach: Insights for the Design of New Antibiotics Drugs.” *Journal of Chemical Information and Modeling* 54 (9). https://doi.org/10.1021/ci5003069.

Silva, José Rogério A, Thavendran Govender, Glenn E M Maguire, Hendrik G Kruger, Jerônimo Lameira, Adrian E Roitberg, and Cláudio Nahum Alves. 2015. “Simulating the Inhibition Reaction of Mycobacterium Tuberculosisl,d-Transpeptidase 2 by Carbapenems.” *Chemical Communications* 51 (63): 12560–62. https://doi.org/10.1039/C5CC03202D.

Woods, R J, and R Chappelle. 2000. “Restrained Electrostatic Potential Atomic Partial Charges for Condensed-Phase Simulations of Carbohydrates.” *Journal of Molecular Structure: THEOCHEM* 527 (1): 149–56. https://doi.org/https://doi.org/10.1016/S0166-1280(00)00487-5.
